# Aversive bimodal associations impact visual and olfactory memory performance in *Drosophila*

**DOI:** 10.1101/2022.07.23.501229

**Authors:** Devasena Thiagarajan, Franziska Eberl, Daniel Veit, Bill S. Hansson, Markus Knaden, Silke Sachse

## Abstract

Insects rely heavily on sampling information from their environment for survival and fitness. Different sensory channels with dedicated downstream neural substrates are programmed to fine tune these signals and translate them into usable instructions to mediate complex behaviors. Sensory information can also be learned and stored as memories that influence decisions in the future. The learning of unimodal sensory signals, especially olfaction, has been studied extensively in different conditioning paradigms in different insects. Using the *Drosophila melanogaster* model in our work, we first investigated differences in the efficiencies of aversive associative visual and olfactory learning using sensory cues that are innately very attractive, such as blue or green light as well as food odors found in fruits or yeast. We then combined the presentation of visual and olfactory sensory stimuli during training to study the effect of bimodal integration on learning performance. When trained unimodally, flies did not easily learn visual stimuli, but when trained bimodally, they developed a significant short-term visual memory after a single learning trial. The bimodal training also suppressed the phototaxis response of the flies to near abolishment. However, a similar training did not enhance the short-term olfactory memory and in some cases, even caused reduction in strength. An enhancement after bimodal training was only seen with a weak long-term olfactory memory retrieved after 24h. Our study demonstrates that bimodal sensory integration is not always synergistic, but is conditional upon the training paradigm and inherent learning abilities of the sensory modalities involved.

## Introduction

All organisms develop intricate sensory relationships with the environment around them. These interactions are essential to sustain and propagate life across diverse habitats. Insects, being one of the most versatile organisms to walk the Earth, depend enormously on these sensory interactions to perform a wide variety of behaviors, therefore providing strong research interests for entomologists and neurobiologists alike. The use of sensory signals to elicit foraging, courtship, mating and predatory behaviors have been long studied and characterized across different insect orders [1–10]. Although insects are equipped with dedicated unimodal sensory apparatus on their bodies and in the brain, signals available in the nature are hardly isolated and insects perform extensive multisensory consolidation before these signals become usable instructions for guiding a behavior. The integration of sensory information deriving from diverse sensory modalities occurs in specific brain centers. In vertebrates, the cortex and the superior colliculus are known to perform this function with the help of bi- and trimodal neurons [11–13]. In insects, different higher-order centers such as the mushroom body, the lateral horn and the central complex are shown to receive multimodal information from different sensory pathways [14–17].

In different scenarios, sensory systems interact with and affect one another. Associations are often made using these sensory signals to remember potential outcomes in a specific situation. Both beneficial and adverse outcomes are learned and retained in the insect brain for either a short or a long period. Research in learning and memory across decades has shown that insects can learn olfactory and visual signals, with some orders showing more acuity in this ability than others. Honeybees can learn to associate odors and several attributes of a visual information such as shapes, colors and patterns with potential rewards or punishments [1,6,18]. Vinegar flies (*Drosophila melanogaster*), while investigated thoroughly for their remarkable olfactory learning ability [19,20], do not exhibit comparable strengths in color learning [21,22]. Ants, on the other hand, learn both olfactory and visual cues in the process of foraging and nest identification but exhibit much faster learning rates, when exposed to bimodal cues [23–25].

Conventional olfactory associative conditioning experiments have been extensively investigated in *D. melanogaster* using a T-maze [20,26,27]. While a specific circular conditioning arena was developed to study color learning in flies [22,28], tethered flight arenas are also commonly used to investigate both pattern learning and multisensory integration during flight behaviors [29–31]. One of the first evidences of cross-modal information transfer was also seen in a flight simulator, where bimodal training of poorly learned unimodal stimuli showed enhanced memory performances in both pattern and odor learning [32]. This observation warrants an important question – is the presence of an additional stimulus reinforcing the same outcome always beneficial in the process of learning or is it conditional upon the requirements of the insect?

In an attempt to answer this question, we aimed to dissect the effects of bimodal associative conditioning on visual and olfactory learning in the fly. Although studies on color vision in *D. melanogaster* have clearly stated the ability of these flies in perceiving and preferring different wavelengths [33–35], work done using color stimuli in associative conditioning experiments have not yielded very high learning performances. In fact, for aversive conditioning assays, flies require intense training trials for achieving high learning scores and the acquired visual memory decays almost entirely after 6 hours [21]. These characteristics of color learning make it a very interesting paradigm to be used as a component of bimodal training. In our work, we adapted and modified the conventional Tully T-maze to present the flies with either unimodal or bimodal olfactory (odors) and visual stimuli (lights of different wavelengths) that are paired with an aversive reinforcement (electric shock). We show that bimodal training selectively enhances weak or reduced unimodal olfactory and visual memories, but the composite bimodal memory itself is not stronger than the unimodal olfactory memory. We therefore propose that bimodal integration has diverse effects on the learning and memory performances of different stimuli.

## RESULTS

### Flies exhibit varied learning performances for olfactory and visual stimuli

In order to investigate the differences in unimodal and bimodal learning of vision and olfaction, we adapted and modified the conventional Tully T-maze **(Fig. 1A)**. Pulsated odor flow and homogeneous illumination from individual LED arrays were paired with the delivery of electric shocks. This design allows for both synchronous and separated presentation of sensory stimuli, depending on the experimental requirements **(Fig. 1B)**. We used two food odors found in fruits or yeast that are attractive for the flies - acetoin acetate (AAC) and ethyl butyrate (ETB) [36] – as well as two wavelengths of light - Blue (452 nm) and Green (520 nm) - that are perceived by distinct visual receptor neurons [28,37,38] and have been shown to be distinguishable in previous visual learning paradigms [22]. We used two kinds of aversive training in our modified T-maze set-up. First, we employed an absolute conditioning paradigm **(Fig. 1C, D)**, where we trained flies to associate an odor or color with an electric shock of 90V. In the testing phase, the flies were allowed to choose between the odor and the solvent (or the illuminated vs. the non-illuminated control arm of the T-maze) in the absence of the reinforcement. As a second learning paradigm we used a differential conditioning protocol **(Fig. 1G, H)**, where we first trained the flies to associate one odor or color (conditioned stimulus, CS+) with the electric shock, while another odor or color was not paired with any reinforcement (CS-). Post training, the choices between the two odors or colors were tested. Naive preference of the flies to the different stimuli was monitored throughout the experiment and an innate preference index (PI) was calculated. The preference indices were used to quantify the change in behavior of the flies towards the odors and the colors before and after training. Each odor and color was trained and indexed separately to assess individual learning capacities.

**Figure 1.**
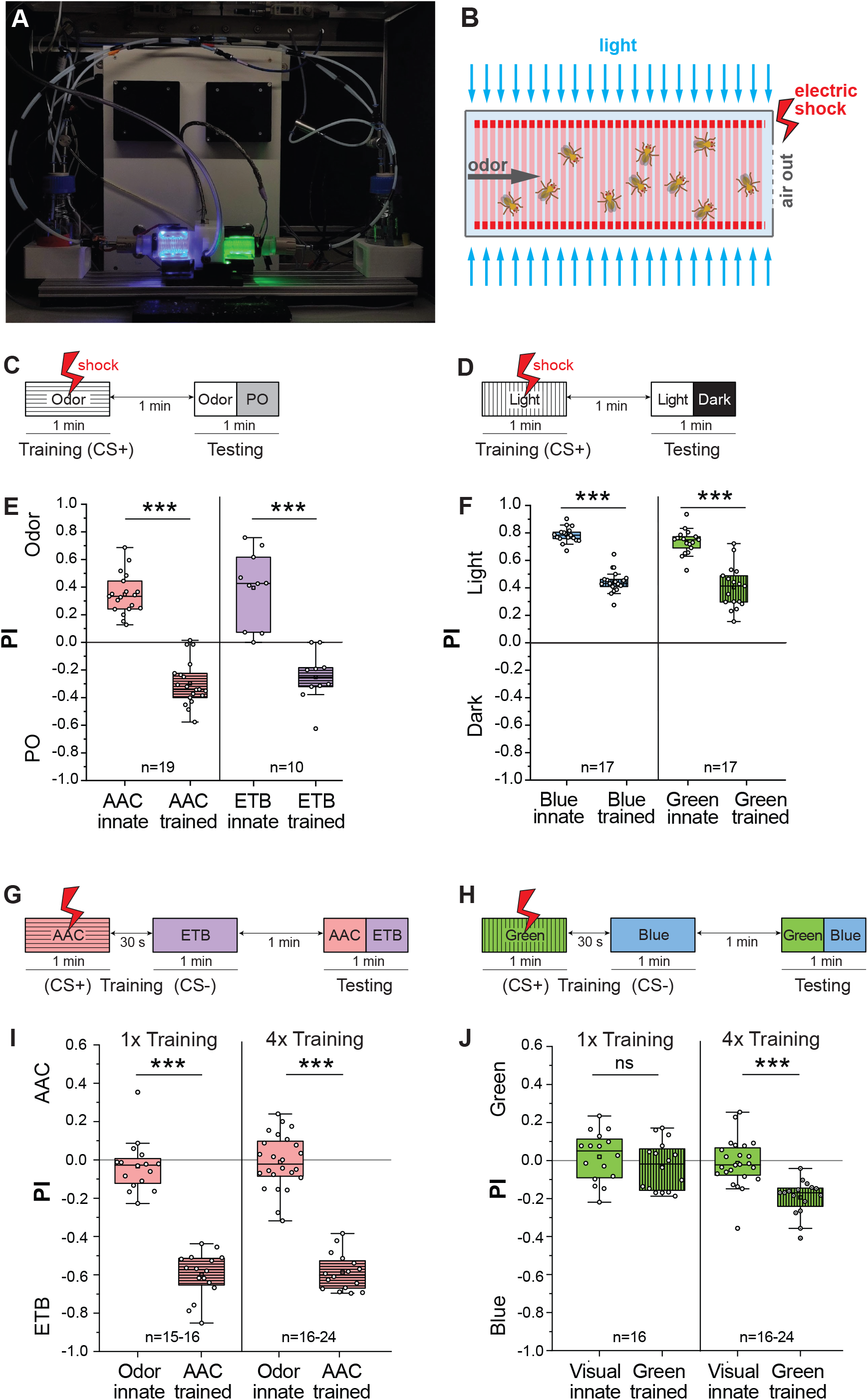
*D. melanogaster* exhibits varied aversive learning responses for different odors and colors. **A**. Image of the T-maze design adapted and optimized from the conventional Tully-T apparatus. The LED contraptions are kept open for illustration purposes. **B**. Schematic of the inside of a training tube during stimuli presentation. **C, D**. Training protocol for absolute olfactory and visual conditioning, respectively. **E**. Preference indices for acetoin acetate (AAC) and ethyl butyrate (ETB) before and after absolute aversive conditioning. Significant aversive olfactory memory was seen in flies for both AAC and ETB. **F**. Preference indices for blue and green wavelengths before and after absolute aversive conditioning. Significant suppression of phototaxis, but no avoidance was seen for both blue and green wavelengths. **G, H**. Training protocol for differential olfactory and visual conditioning, respectively. **I**. Preference indices for AAC before and after one-trial and four-trial (1x/ 4x) training. Aversive olfactory memory was strong and similar between 1x and 4x training protocols. **J**. Preference indices for green before and after 1x/ 4x training. Aversive visual memory in flies is weak and was not seen in the 1x differential training protocol. When training trials are increased to four, a significant visual memory was observed. Each data point represents one experimental trial with 60-80 flies. Comparison between two normally distributed groups were carried out using parametric unpaired two-sample student T-test (***p<0.001).

When presented with a choice against the solvent paraffin oil (PO), the flies showed a significant innate attraction towards the odors. Following one-trial aversive conditioning, this attraction shifted into significant avoidance with the flies choosing the solvent arm consistently **(Fig. 1E)**. In the differentially trained learning paradigm, the concentrations of the odors were standardized beforehand to abolish any innate bias, such that both odors were preferred equally by naïve flies. After differential training, flies exhibited strong aversive learning of the trained odor and shifted their preference to the arm with the CS-odor **(Fig. 1I)**. We also observed that a one-trial differential training was sufficient to achieve the same strength of memory as a four-trial training with an inter-training interval (ITI) of 1 minute emphasizing the ability of the flies to perform very efficient aversive olfactory learning performances **(Fig. 1I)**.

Owing to strong phototaxis behavior, flies showed high preferences to both tested wavelengths when tested for their choice between the illuminated and the non-illuminated arms of the T-maze. In our experiments, we show that unimodal training of the light with an electric shock caused significant reduction in phototaxis **(Fig.1 D, F)**. Unlike previous protocols for visual learning [28], we did not present any electric shock during the test phase in order to arouse the flies. While the reduction of phototaxis indicates a successful reinforcement of the punishment, it is also evident from these results that the strength of the training is not sufficient to entirely overcome the innate phototaxis response of the flies. Conceivably, phototaxis represents a strong innate driving force that cannot be easily countered. Therefore, we again turned to a differential conditioning paradigm to circumvent this issue. Similar to the differential olfactory training, the intensities of the blue and green wavelengths were optimized to ensure minimum innate bias to one of the wavelengths so that naïve flies were equally distributed in both the arms of the T-maze before training. The one-trial differential training of visual stimuli did not yield significant learning performances, but when the training cycle was repeated four times, flies exhibited robust visual learning. **(Fig. 1H, J)**. Put together, our results demonstrate that the same training paradigm results in varying strengths of aversive learning for olfactory and visual stimuli.

### Presence of odor during training enhances visual learning in an absolute conditioning paradigm

After successfully establishing unimodal olfactory and visual aversive conditioning in our set-up, we next aimed to dissect the effect of a bimodal training paradigm on the unimodal visual and olfactory learning performance of flies. We first addressed the question if a combination of two sensory modalities during associative learning can lead to stronger learning efficiency. We delivered three different kinds of CS+, a unimodal olfactory, a unimodal visual and a bimodal (odor + light) stimulus **(Fig. 2A-C)**. Unimodally trained flies were tested only with the trained olfactory or visual stimulus, while bimodally trained flies were tested either with the composite bimodal stimulus or with its individual olfactory and visual components **(Fig. 2C)**. Notably, bimodal training of flies followed by bimodal testing yielded the same strength of memory as unimodal olfactory training **(Fig. 2D)**. However, it is difficult to conclude the individual contribution of each sensory modality in this composite memory performance, especially since an additive effect was not evident. We therefore turned our attention to the individual retrieval of olfactory and visual memory responses after bimodal training. The olfactory memory of bimodally trained flies was not significantly different from that of the flies that were trained unimodally. We hypothesize that this lack of enhancement could be attributed to the already strong olfactory memory, which cannot be further enhanced by the addition of a visual stimulus during training. Interestingly, when flies were trained bimodally but tested unimodally with only the light stimulus, we observed a striking reduction in the phototaxis response. Hence, an improved visual memory was observed in comparison to the unimodally trained to the visual stimulus. In this experiment, the absence of the odor during the test phase negates any direct effect of the strongly learned odor response and retains only the visual memory following bimodal training. These results indicate that the presence of an odor along with a light stimulus leads to stronger acquisition of a visual memory.

**Figure 2.**
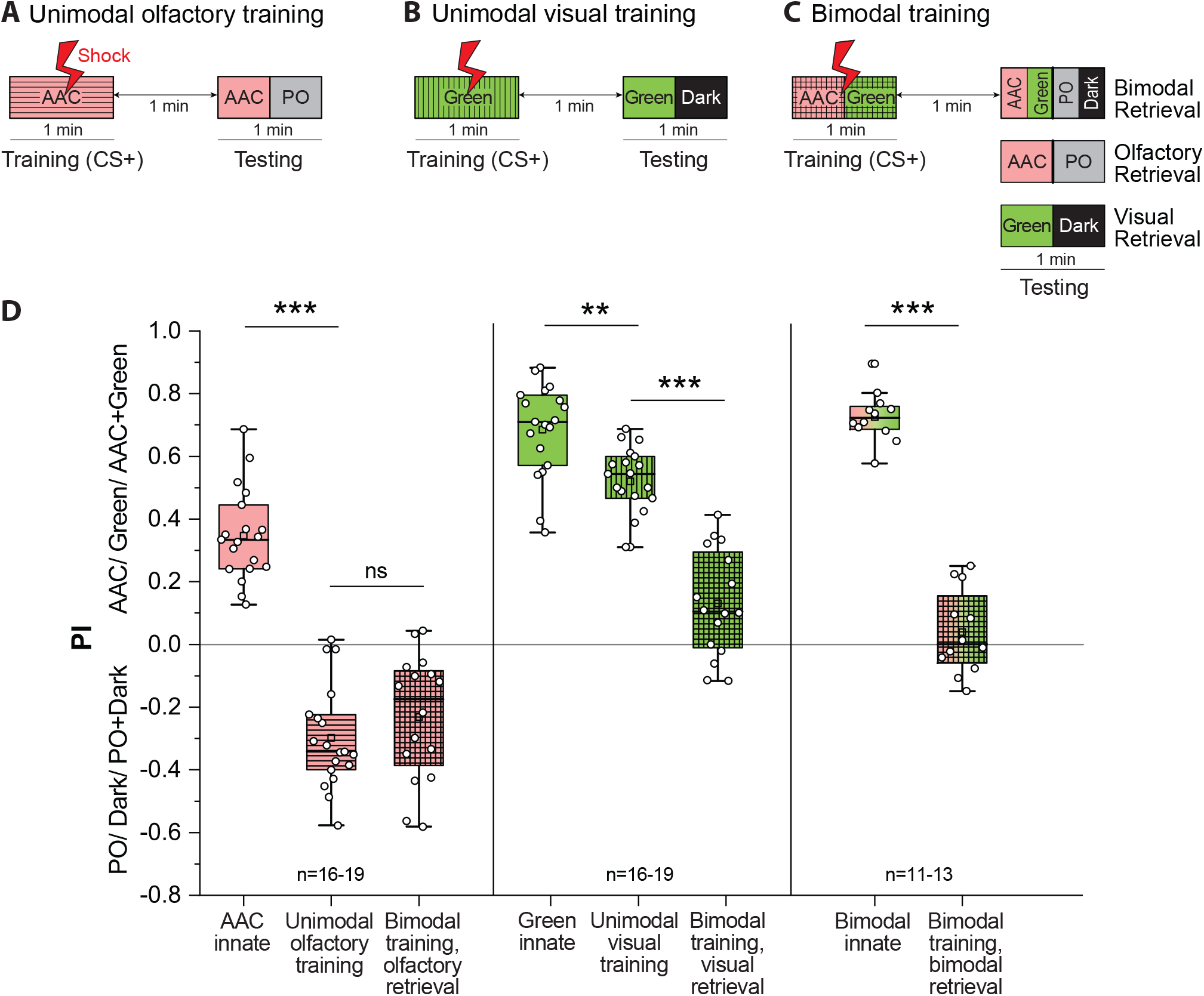
Presence of odor during absolute conditioning enhances unimodal visual memory. **A, B**. Training protocol for absolute aversive unimodal olfactory and visual conditioning. **C**. Training protocol for absolute aversive bimodal conditioning with three different testing protocols. **D**. Preference indices for AAC, Green and AAC and Green plotted before and after unimodal and bimodal training. Visual memory retrieval after bimodal training shows much stronger phototaxis compared to only unimodal visual training. Olfactory memory retrieval after bimodal training does not differ significantly from only unimodal olfactory training. Statistical comparisons between more than three normally distributed data sets were made using one-way ANOVA followed by Bonferroni correction and Tukey’s post-hoc test (**p<0.01, ***p<0.0001). Comparison between two normally distributed data sets was made using unpaired Student T-test with a Welch’s correction (***p<0.0001)

### One-trial differential bimodal training of colors yields significant visual memory

We next turned to the differential training paradigm to test whether a similar enhancement of visual memory also occurs in another learning paradigm **(Fig. 3A)** Innate preferences were always tested for all experimental conditions before training and they served as controls for comparison with the learned responses. Bimodal training consisted of one odor and color combination (e.g. AAC and Green) serving as CS+ and another combination serving as CS-(e.g. ETB and Blue). Following bimodal training, testing was carried out in three different categories as described before to analyze (i) a composite bimodal memory, (ii) an olfactory memory and (iii) a visual memory. We performed a one-trial differential bimodal training (i.e. 1x) for all possible odor and color combinations. For deducing the effect of bimodal training only on the visual memory, we trained the flies to the bimodal stimulus but tested them afterwards only to the light stimulus, excluding the effect of a learned odor bias **(Fig. 3A)**. We observed that a significant avoidance was displayed towards the trained color even in the absence of the paired odor during testing. The addition of odor information during the training procedure induced a significant visual memory to emerge, while training with only a unimodal visual stimulus did not yield any memory. This effect was evident across all color and odor combinations **(Fig. 3B-E)**. Conversely, in order to identify the effect of bimodal training on only the olfactory memory, we trained the flies to the bimodal stimulus but tested them afterwards only to the odor stimulus, excluding the effect of any color bias. Notably, in these experiments we observed a reduction in the strength of the olfactory memory. In two combinations (AAC+Blue and ETB+Blue), the unimodally retrieved olfactory memory after bimodal training was significantly weaker than the unimodal olfactory memory **(Fig. 3D, E)**, while we observed a trend of reduction in the other two combinations. In three combinations, the composite bimodal memory was weaker when compared to the unimodal olfactory memory **(Fig. 3C, D, and E)**.

In order to confirm the role of the olfactory input in enhancing a short-term visual memory, we performed the same experiments for one combination (AAC and Green) with olfactory mutant flies that lack the odorant receptor co-receptor (*Orco*) and the ionotropic receptor co-receptor (*Ir8a*) **(Fig. 4A)**. While *Orco* is coexpressed with odorant receptors (ORs) and is crucial for proper OR functioning, *IR8a* plays a role in IR-mediated acid and aldehyde sensing [39]. Mutant flies lacking the expression of *Orco* and *Ir8a* cannot detect the odors that were used in our assay and therefore did not show any significant olfactory memory. Subsequently, they also did not reveal any significant visual memory after being bimodally trained to both an odor and a visual stimulus **(Fig. 4B)**. These results emphasize the importance of an olfactory stimulus, especially the learned response to the odors, in the formation of a visual memory after bimodal training. This observation further confirms that bimodal integration underlies the enhancement of visual memory.

**Figure 3.**
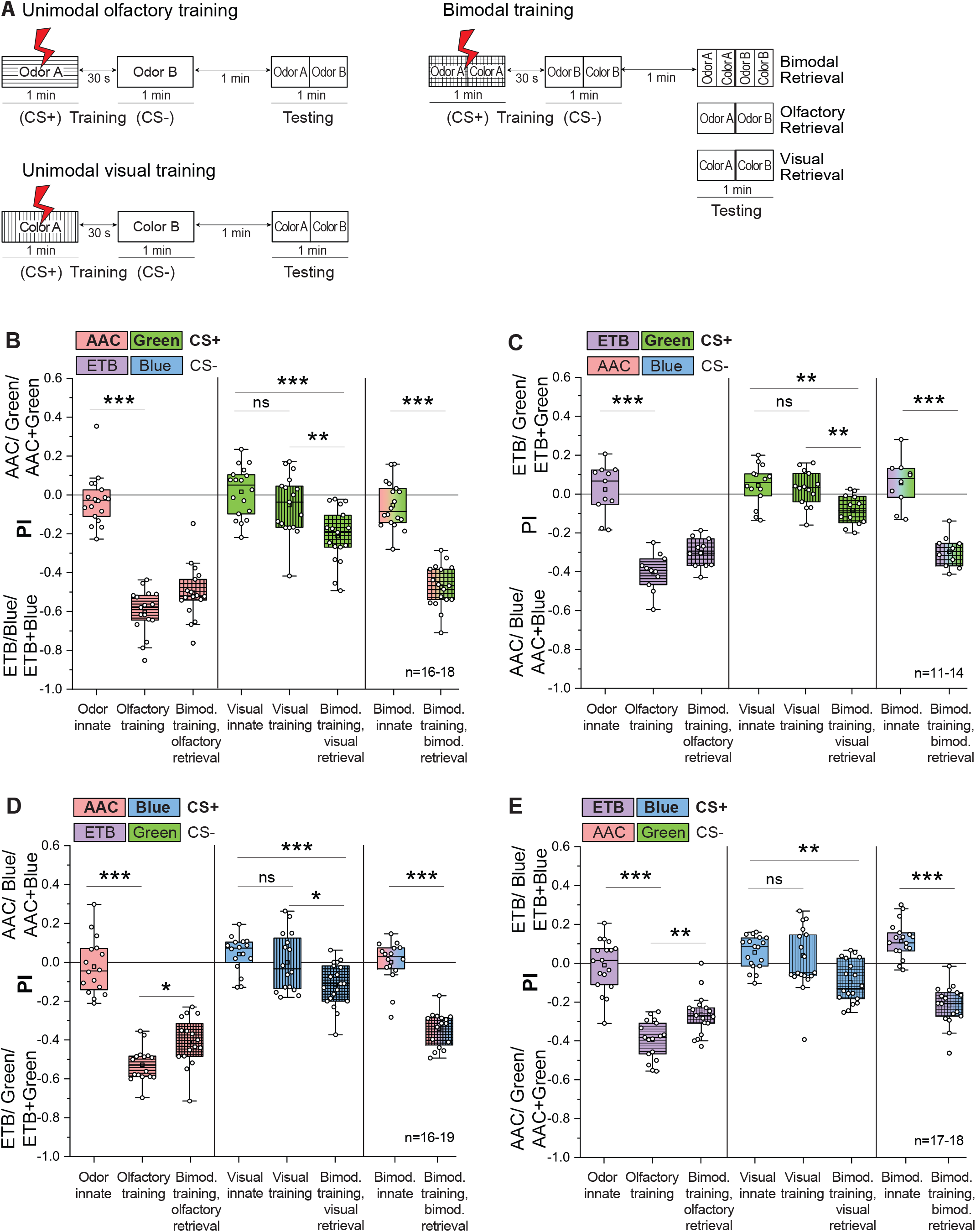
One-trial differential bimodal training of colors yields significant short-term visual memory. **A**. General training protocols for differential unimodal olfactory, visual and bimodal training along with corresponding testing procedure. **B, C, D, E**. Preference indices before and after unimodal and bimodal differential training, plotted for different combinations of odors and colors. Emergence of visual memory can be seen across combinations after bimodal training. A reduction of olfactory memory was also observed when flies were bimodally trained, but unimodally retrieved. Composite bimodal memory was not stronger than unimodal olfactory memory. All statistical comparisons between normally distributed data sets were made using one-way ANOVA followed by Bonferroni correction and Tukey’s post-hoc test. Kruskal Wallis test followed by Dunn-Bonferroni correction was done for comparisons between three or more groups that had non-parametric properties (*p<0.05, **p<0.01, ***p<0.001).

**Figure 4.**
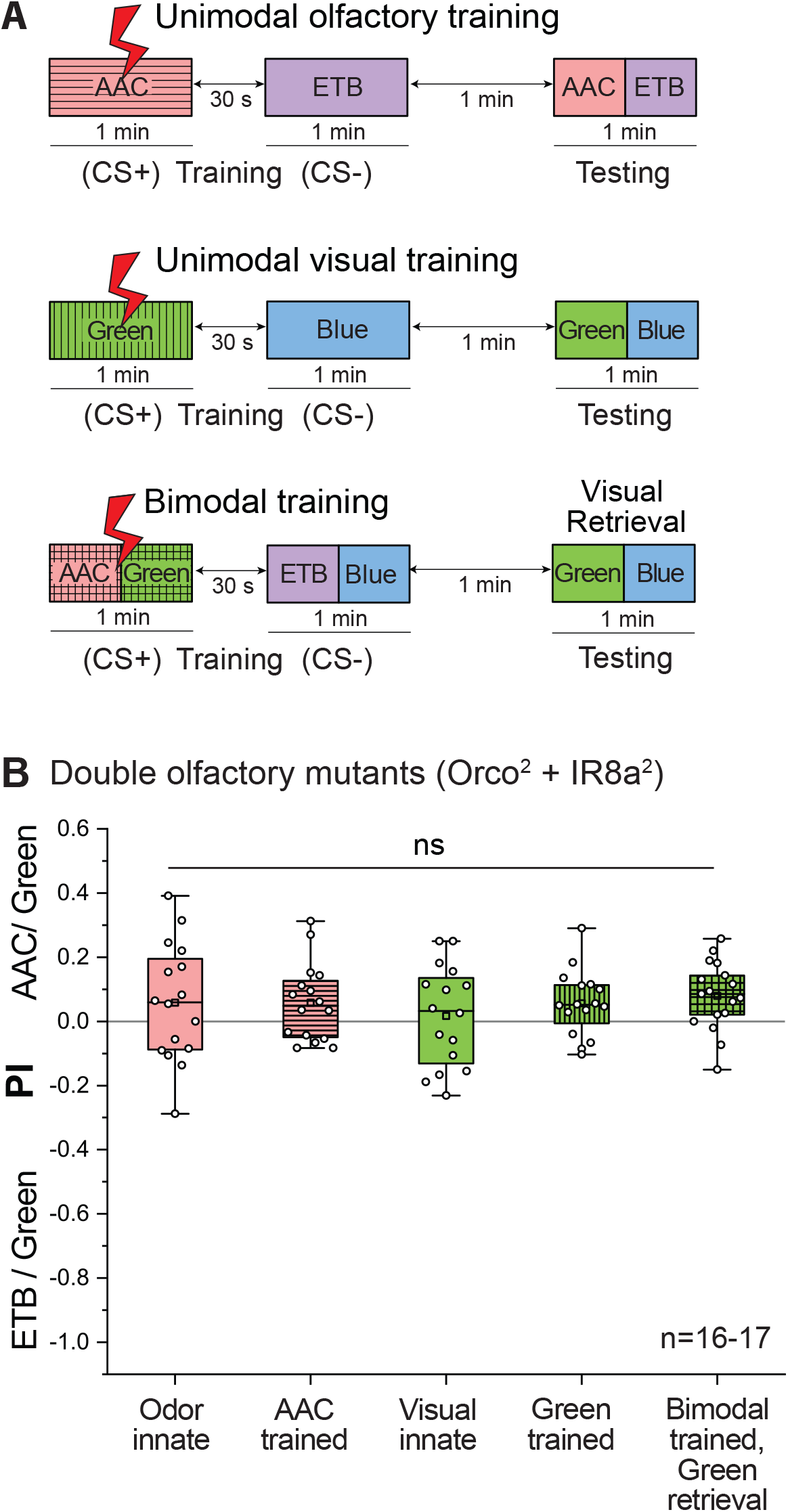
Olfactory mutants do not show an enhanced short-term visual memory after bimodal training. **A**. General training protocols for differential unimodal olfactory, visual and bimodal training along with corresponding testing procedure. **B**. Preference indices before and after unimodal and bimodal differential training for double olfactory mutants lacking *Orco* and *IR8a* co-receptors. Inability of the flies to detect odors abolishes olfactory learning and therefore does not show a significant visual memory after bimodal training. All statistical comparisons were made using one-way ANOVA followed by Bonferroni correction and Tukey’s post-hoc test (ns – not significant, p>0.05)

### Bimodal training selectively enhances weakened olfactory and visual memories

Our initial experiments showed that flies exhibit a significant visual memory in a differential conditioning paradigm after four training trials **(Fig. 1J)**. In order to investigate whether the strength of the observed visual memory can be enhanced even further, we performed four-trial bimodal training experiments (i.e. 4x) with an inter-trial interval of 1 minute using one odor-color combination (AAC and Green) **(Fig. 5A)**. Although we observed a trend of an increase when we trained the flies to a bimodal stimulus and retrieved only the visual memory, it was not significantly greater than unimodal visual memory, indicating that an upper limit might exist for the learning strength of visual stimuli. Additionally, similar to the previous experiments (**Fig. 3D, E**), a significant decrease was observed in the strength of the olfactory memory when it was retrieved separately after four trials of differential bimodal training **(Fig. 5B)**. We also note here that the composite bimodal memory retrieved after bimodal training was not stronger, but rather similar to the unimodal olfactory memory.

**Figure 5.**
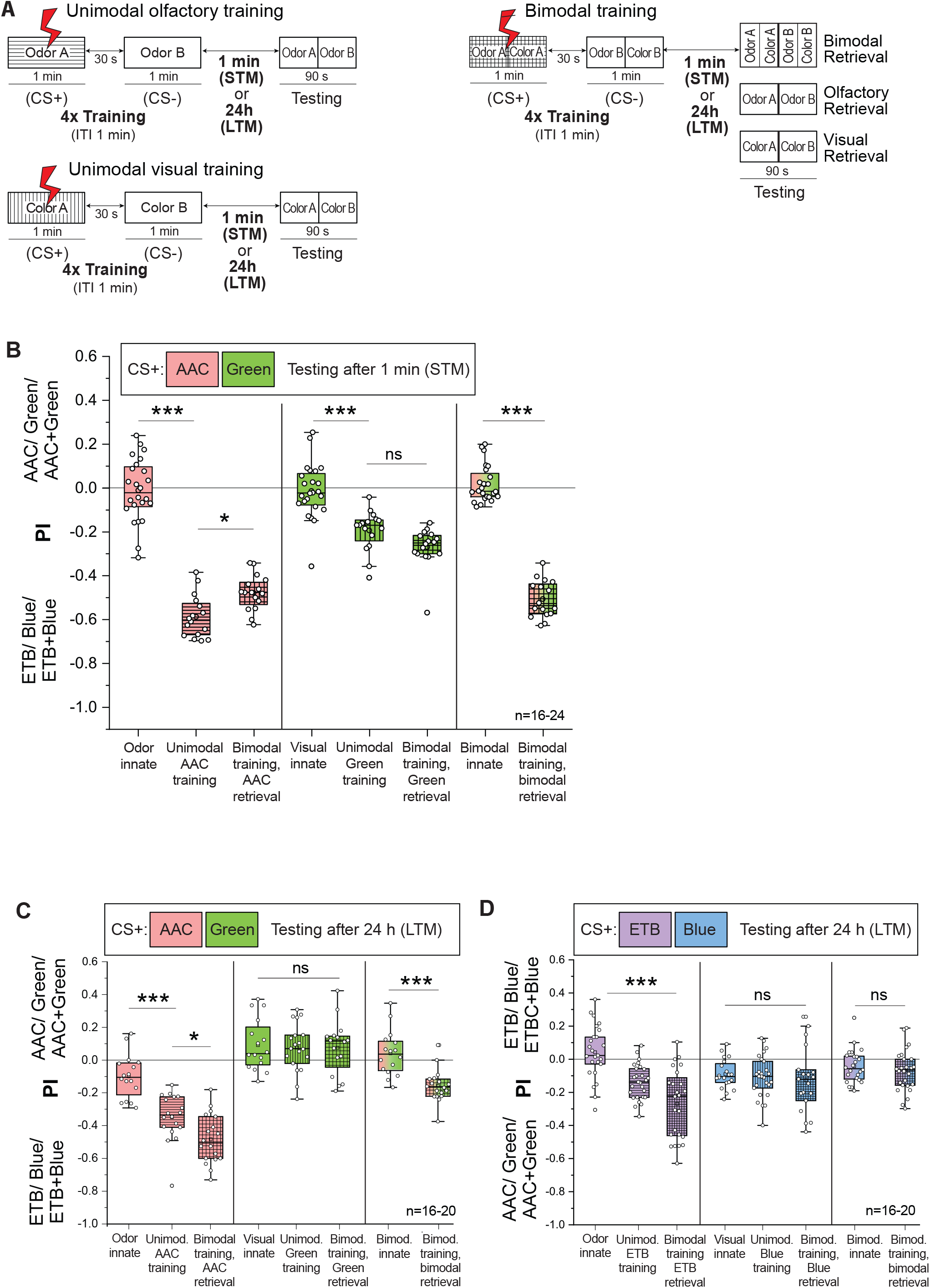
Bimodal training selectively enhances weakened olfactory and visual memories. **A**. General four-trial (4x) training protocols, with an inter-training interval of 1 min, for differential unimodal olfactory, visual and bimodal training along with corresponding testing protocols. Short-term testing was done 1 min after the last training cycle. Long-term testing was done 24 h after the last training cycle. **B**. Short-term memories retrieved after four-trial training plotted as changes in preference indices of the flies to AAC, Green and AAC + Green before and after unimodal/ bimodal training. Four training trials lead to significant short-term visual memory to form. This is not enhanced further when bimodally trained, but unimodally retrieved. A reduction of olfactory memory was observed when bimodal trained, but unimodally retrieved. Composite bimodal memory was similar in strength to unimodal olfactory memory. For comparisons between groups with odor treatment, one-way ANOVA followed by Bonferroni correction and Tukey’s post-hoc test was done (*p<0.05, ***p<0.001). For comparisons between groups with visual treatment, Kruskal Wallis test followed by Dunn-Bonferroni correction was done (***p<0.001). **C, D**. Long-term memories (LTM) retrieved after four-trial training plotted as changes in preference indices of the flies to different combinations of stimuli, before and after unimodal/ bimodal training. Strengthening of olfactory LTM was observed following bimodal training. Complete absence of visual LTM was also observed. For comparisons between normally distributed datasets, one-way ANOVA followed by Bonferroni correction and Tukey’s post-hoc test was done (*p<0.05, ***p<0.001). For comparisons between groups with visual treatment, Kruskal Wallis test followed by Dunn-Bonferroni correction was done (***p<0.001).

In experiments where testing immediately followed the one-trial training protocol, we observed that bimodal training had an incremental effect on weak/ non-existent visual learning. These results provide evidence for a learning enhancement in at least one modality as a consequence of bimodal integration. However, the observation that we could not achieve a similar effect for olfactory learning led us to investigate potential underlying causes. We hypothesized that, while retrieved as a short-term memory (STM), both one- and four-trial aversive olfactory training protocols achieved maximum learning efficiency, with little capacity for further memory enhancement. In addition, we also considered that enhancement following bimodal integration, if any, might not be prominently visible in our assay with such high unimodal learning scores. In order to test these hypotheses, we worked with a compromised or weakened olfactory memory that would provide us a window to observe an improvement. We performed experiments using the four-trial differential training protocol described before, but tested for long-term memory (LTM) after a period of 24 h **(Fig. 5A)**. Innate preferences to different stimuli were observed and indexed on the day of testing. Two different combinations were used as CS+ (i.e. AAC and Green or ETB and Blue).

As expected, unimodal LTM for both odors was weaker than the STM that was obtained after the four-trial differential training protocol, with ETB retaining an even lower LTM than AAC **(Fig.5C, D)**. Notably, when bimodally trained but retrieved only with the odor after 24 h, flies exhibited a strengthened long-term olfactory memory for the odors. However, consistent with previous studies, we observed no visual LTM. This led us to conclude that in spite of not retaining any significant LTM of its own, simply the presence of a visual stimulus during the training procedure leads to a stronger long-term olfactory memory. We also observed a reduced composite bimodal memory in case of AAC and Green and none for ETB and Blue when compared to the unimodal olfactory learning or short term bimodal learning. We attribute this observation to the complete lack of a visual LTM and the weakly retained LTMs of the respective odors.

## DISCUSSION

We developed a modified T-maze that was adapted from the conventional Tully-T to perform aversive unimodal and bimodal learning experiments. Our set-up consisted of simplified T-arms and the use of an improved odor delivery system that ensured consistent odor concentration throughout the experimental duration (observed and quantified for up to 6 h). We used a pulsed odor delivery that was synchronized exactly to the shock presentation, providing maximum overlap between them (see Materials and Methods). In addition, we employed LED panels to deliver the light stimulation in a homogeneous manner along the length of our training and testing tube. The use of the same experimental apparatus to study both aversive olfactory and visual learning aided in studying the differences in learning efficiencies of the two sensory modalities. Using this method, significant learning scores were achieved after aversive visual training. Furthermore, the odors that we used in our study are attractive food-derived compounds that elicit innate attraction in flies. The high learning scores that were observed for these odors and colors reaffirm the versatility of our assay.

### Learning and memory performances vary for distinct stimuli, both within and across modalities

In our study, we demonstrate that identical training paradigms yield varying learning performances for different odors and colors. *D. melanogaster* exhibits a very strong short-term aversive olfactory memory following a one-trial training paradigm. We show that repeating the same training cycle for four times with an ITl of 1 minute does not lead to an enhancement of learning. However, the same one-trial training paradigm did not generate any visual learning, which required repeated reinforcements with four training trials. This could be attributed to the large extent up to which flies rely on olfaction to perform diverse behaviors. It is a clear advantage to learn and remember associations, which are essential for survival in nature using the most reliable sensory information that is also perceived with high precision by the insect. Although shared circuits in the mushroom body underlie both visual and olfactory learning [22], with our experiments, we also see differences in the ease of acquisition, strength of learning and retention between the two modalities, suggesting that intricate differences may still exist in the learning circuitry for both of them. The significant suppression of phototaxis after one-trial absolute conditioning can still be regarded as clear visual learning, while the same training protocol in a differential paradigm did not yield similar results. In the latter paradigm, flies required the training cycle to be repeated four times to show a significant aversive visual memory. It can be speculated that wavelength discrimination as a consequence of conditioning requires intense training trials, while the discrimination between the presence and absence of light (as in the case of absolute conditioning) can be learned and remembered more easily. These results indicate that unimodal visual learning, specifically discriminatory color learning, may not be employed by *D. melanogaster* in nature as extensively as odor learning to mediate crucial behavioral decisions.

### Bimodal integration has opposing effects on olfactory and visual memory performances

Research involving bimodal sensory processing in the past has proven that cross-modal facilitations enhance innate as well as memory performances of the interacting sensory modalities [40–44]. However, our work shows that such enhancements in the context of learning are conditional. Using wavelength discrimination as form of weak visual learning, we corroborate the work done in the tethered flight arena, where sub-threshold pattern learning of heat beams was enhanced by the addition of odor during the training [32,45]. Furthermore, we also provide evidence that olfactory stimuli that are already strongly learned and memorized, do not benefit from bimodal training. Our results even indicate a reduction in the short-term olfactory memory when retrieved separately after bimodal training, because it might be negatively impacted by the presence of a weakly learned visual stimulus **(Fig. 3B-E)**. Weak visual memory showed a clear improvement after bimodal training specifically in experiments where unimodal training was fruitless **(Fig. 3B-E)**. However, we also demonstrate that in a four-trial associative training, where an upper capacity for unimodal visual memory is already achieved, a significant enhancement could no longer be seen even after bimodal training **(Fig. 5B)**. We also show an increase in the reduced olfactory LTM after bimodal training. This could be either an enhancement effect or a stabilization in the LTM circuit that slows down the decay of the memory trace and improves memory retrieval. However, this mechanism is difficult to pinpoint without further experiments targeting the memory consolidation process after unimodal and bimodal training. When using weakly learned stimuli such ETB or Blue, we were not able to see a bimodal memory after 24 h. This implies that the inherent strength of unimodal memories could also contribute to the extent up to which bimodal training can affect them.

Learned outcomes in nature that cannot be associated or retrieved by one sensory modality can be supplemented by the presence of a different, strongly associated sensory modality. However, when the reinforcements are strong enough to sustain a unimodal memory association, the presence of an additional modality does not augment the learning anymore and can even cause detrimental effects if the additional modality is in itself weakly learned. In conclusion, we see that the effect of bimodal integration is not always synergistic or additive, but is conditional upon the learning abilities of the participating modalities. Future studies should be dedicated to elucidate the underlying neuronal circuitry and to pinpoint the neuronal sites in the fly brain accomplishing multimodal integration. While visually-selective Kenyon Cells in the mushroom bodies are required for appetitive bimodal associative learning as recently shown [46], the neuronal substrate for aversive bimodal associative learning as used in our study most likely represents distinct circuits which remain so far elusive.

## MATERIALS AND METHODS

### Fly rearing

Corn-meal agar was used as a food medium to raise flies at 25°C and 70% humidity in incubators manufactured by SnijdersLabs. 12:12 Light/ Dark cycle was followed to preserve circadian rhythms. All behavioral experiments were performed with wild-type Canton-S flies, except in one experiment with olfactory mutants where a double Orco2 / Ir8a2 mutant was used. This line was a kind gift from Yael Grosjean. Adults of both female and male sexes were used in the learning assays.

### Stimulus presentation in behavioral experiments

#### a. Odor delivery

Two attractive odors were used in our experiments - acetoin acetate (AAC, CAS No. 4906-24-5, SIGMA) and ethyl butyrate (ETB, CAS No.105-54-4, SIGMA). As we observed highly fluctuating odor concentrations during the span of the experiment with the conventionally used cubical contraptions, we adapted a new method of odor delivery. Custom-modified 4 mL HPLC glass vials (supplier) containing 1 mL of odorant were fitted with a metal capillary of a maximum diameter of 2.5 mm and were placed inside larger 250 mL bottles (Schott GmBH). Clean, compressed and humidified air was allowed to pass through these bottles at a constant and controlled flow rate of 0.35 L/min, which was then carried along with the headspace of the bottle to the T-maze using Teflon tubes. Flow rates were controlled using digital flow meters. Volatile odor molecules in the glass vials evaporated into the headspace. An equilibrium of odor concentration was established and maintained with the headspace in the larger bottles. Standardization of the volatile measurement was done using Gas Chromatography coupled Mass Spectrometry (GC-MS). We observed that one hour of odor accumulation in the larger bottle followed by one hour of flushing with clean air was required to achieve consistent odor concentrations for a total pulsing duration of 6 h using the pure AAC and 1% ETB (solvent – paraffin oil, PO) in the glass vials. These concentrations elicited equal preference from the flies in a choice assay. Odor at a flow rate 0.35 mL/minute was delivered in pulses of 2 s followed by 2 s of only humidified air with 72% relative humidity maintained in the entire arena. Odor pulses were controlled with fast solenoid valves (Festo – Esslingen, Germany).

#### b. Light delivery

Custom made (3D-printed) contraptions with non-reflective black coating on the outside and a fitted LED panel on the inside (Roschwege, Greifenstein, Germany) of specific wavelengths - Blue (452 nm) and Green (520 nm) - were used to surround the entire length of the testing and the training tubes to ensure homogenous illumination of the arena. The training and testing tubes were made of transparent polymethyl methacrylate (PMMA). An intensity dimmer (HK Datentechnik, Dormagen, Germany) was fitted with the power supply for the LEDs (Meanwell, Taiwan) to standardize the optimal light intensities for the two wavelengths that ensured neutral innate preferences for the flies when given a choice.

#### c. Synchronization with the electric shock

The synchronization of the light stimulus, odor pulses and the electric shock was programmed using LOGO Direct Digital Control (DDC - Siemens LOGO, Munich, Germany). The odor pulses and the electric shock were synchronized to ensure sufficient temporal overlap and the light was presented uniformly across the duration of the protocol. A delay of approximately 200ms was measured in the odor delivery and was taken into account, while programming the synchronization.

### Aversive Conditioning Protocol

The T-maze was a custom-made 3D printed arena (white or black polyoxymethylene material) adapted and simplified from the conventional Tully T-maze. For experiments with phototaxis, black arenas were used to ensure least interference from the reflected light during the testing phase. A group of 70-80 flies that were 4-5 days old were aspirated into the training tube that was lined with an embedded copper grid (CON-Elektronik, Germany). Aversive reinforcement of the stimulus (CS+) was provided for one minute (odor/ light/ odor + light presentation) paired with 90V electric shock. Training consisted of 15 shock pulses, each pulse lasting 1.8s followed by an interval of 2.2 s. After training, flies were given 30 s of clean air. In case of absolute conditioning, this interval was for 1 min and then followed by one minute of testing (odor vs paraffin oil). In differential conditioning, the 30 s interval was followed by 1 min of CS-presentation (without shock). One minute after the presentation of CS-, testing was done for 1 min or 90s in some experiments. Suction was always provided to prevent odor contamination or accumulation in the center of the maze. The tube lined with the copper grid was used in both the training and the testing phase to reproduce the same contexts. All training and testing were done at 25°C in a dark arena. Flies inside the arena were handled only in red light. Preference indices were calculated for the innate choice as well as the learned choices using the following formula:

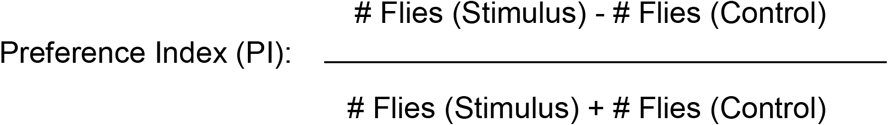

In experiments where long-term memory retrieval was necessary, trained flies were collected and stored in vials with food at 25°C and darkness for 24 h.

### Statistical Analysis

All statistical analyses and graphs were made using the software OriginPro2019 9.6.0.172 (OriginLab corporation). At a significance level of 0.05, data sets were checked for normal distribution using the Shapiro-Wilk test. When normality was established, comparisons between the groups were done using One-Way ANOVA followed by either Bonferroni correction or a post-hoc Tukey’s test. The significance level was set at 0.05. For data with non-parametric distribution, Kruskal Wallis test and a post-hoc Dunn-Bonferroni test were carried out. Comparison between two normally distributed groups was done using unpaired two-tailed Student’s T-test.

## Author contributions

Research design - S.S, D.T, M.K. Experimental work and data collection - D.T, D.V, F.E. Data Analysis - D.T, F.E. Writing – D.T, S.S Revision – D.T, S.S, F.E, M.K, B.S.H. Supervision S.S.,M.K. Funding Acquisition S.S., B.S.H, D.T

## Acknowledgements

**T**he Max Planck Society (MPG) and the Deutsche Akademsicher Austauschdienst (DAAD – stipend to D.T) funded this work. We thank Silke Trautheim and Roland Spiess for their excellent support in fly rearing. We also thank Saskia Gablenz (preliminary experimental design), Florencia Campetella and Veit Grabe (scientific advice and assistance in learning experiments) and Felix Feistel (GC-MS data acquisition and analysis). We finally thank Swetlana Laubrich for her outstanding administrative support.

## Notes

### Competing Interest Statement

The authors have declared no competing interest.

